# Molecular plasticity to soil water deficit differs between sessile oak (*Quercus Petraea* (Matt.) Liebl.) high- and low-water use efficiency genotypes

**DOI:** 10.1101/2021.09.16.460634

**Authors:** Gregoire Le Provost, Theo Gerardin, Christophe Plomion, Oliver Brendel

## Abstract

**Background:** Water use efficiency (WUE) is an important adaptive trait for soil water deficit. The molecular and physiological bases of WUE regulation in crops have been studied in detail in the context of plant breeding. Knowledge for most forest tree species lags behind, despite the need to identify populations or genotypes able to cope with the longer, more intense drought periods likely to result from climate warming.

**Results:** We aimed to bridge this gap in knowledge for sessile oak (*Quercus Petraeae* Matt. L.), one of the most ecologically and economically important tree species in Europe, using a factorial design including two genotypes (low and high WUE) and two watering regimes (control and drought). By monitoring the ecophysiological response, we were able to identify groups of genotypes with high and low WUE. We then performed RNA-seq to quantify gene expression for the most extreme genotypes exposed to two watering regimes. By analyzing the interaction term, we were able to capture the molecular strategy of each group of plants for coping with drought. Regardless of water availability, the high WUE genotypes overexpressed genes associated with drought responses, and the control of stomatal density and distribution, and displayed a downregulation of genes associated with early stomatal closure and high transpiration rate. High-WUE genotypes, thus, coped with drought by fine-tuning the expression of genes with known functions in the regulation of stomatal size, density, movement or aperture and transpiration rate.

**Conclusion:** Fine physiological screening of sessile oaks with contrasting WUE, and their molecular characterization i) highlighted subtle differences in transcription between low and high WUE genotypes, identifying key molecular players in the genetic control of this trait, and ii) revealed the genes underlying the molecular strategy that had evolved in each group to cope with water deficit, providing new insight into the value of WUE for adaptation to drought.

## Background

Maintenance of the net primary production of trees, and, therefore, of the capacity of planted and natural forests to buffer the effects of global warming, is dependent on various biotic and abiotic factors, including their ability to adapt to water shortages during the growing period [1].

It has been suggested that water use efficiency (WUE), defined as biomass production divided by the water used through transpiration [2, 3], plays a functional role in adaptation to water scarcity. Indeed, this trait, whether assessed instantaneously or through integrated measurements, is determined environmentally (as a function of soil water deficit, vapor pressure deficit or temperature) and genetically (reviewed in [3]). Maritime pine, a tree species scattered throughout the Mediterranean basin, constitutes a unique case study illustrating these findings. The phenotypic plasticity of WUE has been shown to depend on soil water deficit and evaporative demand [5, 6], and ample genetic variation has been found both within breeding populations [5, 7] and between natural populations adapted to local climatic or edaphic selection pressures [8–10]. One study [11] also reported a significant GxE interaction, suggesting that maritime pine gene pools have adopted different WUE strategies in response to drought.

Sessile oak (*Quercus petraea* (Matt.) Liebl.)) is a dominant species in Europe. It has a very wide ecological niche, tolerating pH values from 3.5 to 9; it can grow tolerate all degrees of soil wetness, from xeric to wet soils, and it is present in a zone extending from southern Spain to Scandinavia, with a large climatic amplitude [12]. It is relatively tolerant to drought and poor soils, but is sensitive to highly hydromorphic conditions. It has been reported to have a variable WUE directly linked to stomatal conductance [13]. Both environmental (plasticity to drought) and genetic variation (within and between populations) have been shown to drive this variation [13–15].

Studies in oaks have contributed to our understanding of the physiological mechanisms underlying WUE diversity in trees. Studies in pedunculate oak genotypes presenting extreme phenotypic values for WUE [14, 16] clearly demonstrated in well-watered conditions the genetic effect of stomatal conductance on leaf and whole plant water use efficiency, particularly in terms of stomatal density and the diurnal dynamics of stomata.

Several studies have investigated the molecular mechanisms underlying WUE variation in annual. Most of these studies were conducted in the model plants *Arabidopsis thaliana* and *Oryza sativa* (reviewed in [17]). Changes in stomatal aperture have been shown to be linked to both drought stress tolerance and WUE [18, 19]. Based on these findings, several genes involved in the stomatal development pathway have emerged as promising candidate genes for the control of WUE (reviewed in [20]). In particular, the transcription factor GTL1 has been shown to regulate WUE by modifying stomatal density through repression of the SDD1 gene [20]. The HARDY gene, encoding a AP2/ERF-like transcription factor, has been reported to affect WUE directly in rice, by enhancing photosynthetic assimilation and efficiency [21]. In *Arabidopsis*, the *ERECTA* gene, encoding a putative leucine-rich receptor-like protein kinase (LRR-RLK) regulates WUE through its action on leaf morphology (i.e. mesophyll proliferation) and stomatal density [22]. Other authors have reported that the AtEDT1/HDG11 gene, encoding a Homeodomain START (HD-START) transcription factor, also regulates stomatal density and WUE through its interaction with the ERECTA and E2Fa in both *Arabidopsis* and tobacco [reviewed in 18]. In maize, the Asr1 transcription factor regulates WUE by modulating C4 (C4-PEPC) activity [24]. A possible role for carbonic anhydrase, which catalyzes CO_2_ hydration, has also recently been suggested. Indeed, the WUE of double mutants for anhydrase carbonic genes is affected in drought stress conditions, suggesting an important role for these genes in maize [25]. The molecular mechanisms involved in WUE regulation in forest trees have been little studied, and we are aware of only two studies, one in oak [14] and the other in poplar [26], confirming the regulation of WUE by known candidate genes (ERECTA and the GTL1) identified in studies on annual plants.

In this study, we aimed to bridge this gap in our knowledge, by investigating molecular signatures in the leaves of sessile oak genotypes with extreme phenotypic values for WUE subjected to drought. We used a full factorial design in which oak seedlings with contrasting WUE (G) were raised in drought and control conditions (E) and characterized phenotypically and molecularly. Transcripts presenting a E or G effect can be used to identify important gene pathways responding to drought and underlying WUE variation, respectively. We were able to assess adaptive function from the analysis of genes presenting GxE interactions, revealing molecular strategies based on high-WUE genotypes in an environment in which water supply is limited. This study provides important clues to the adaptive value of WUE in sessile oak and the underlying molecular mechanisms. The genes identified here represent new tools to help foresters to identify the most drought-resilient populations or individuals in silvicultural practice.

## Results

### Ecophysiological variation

A Principal components Analysis (PCA) was performed for the four traits used for selection (TE, TE_d_, Wi_in situ_,Wi_SS_) and also for all traits that were measured (N=18). For the four-trait PCA, the first axis explained 83.5%, and the second axis 12,4% of the observed variation. The first axis separates clearly the chosen genotypes in terms of WUE (Figure 1a), whereas the first and the second axis (upwards sloping diagonal) separates the drought groups. For the 18-trait PCA (Figure 1b), the first axis represented 51.9% and the second axis 15.9% of the observed variation. The first axis is strongly related to measurements of WUE (including δ ^13^C), but also stomatal conductance measurements. The separation of drought groups (upwards sloping diagonal) corresponds mainly to whole plant traits such as CT and BM.

**Figure 1:**
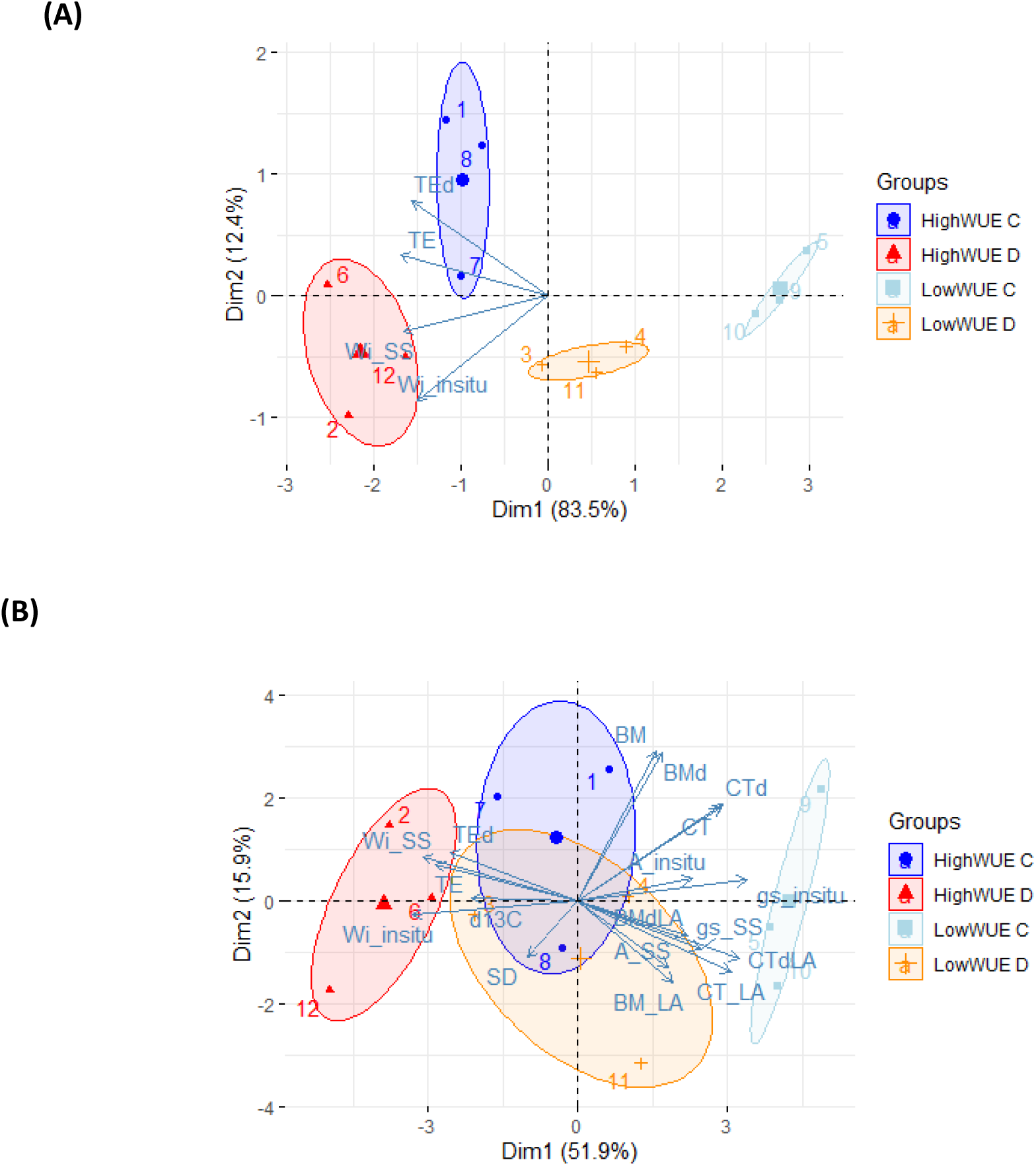
Biplot (individuals and variable vectors) of the PCA analysis of a) the four traits used for genotype selection (TE, TEd, Wiin situ,WiSS) and b) all of the traits presented in Table 1. The four groups are represented as confidence ellipses around the group mean.

#### Drought stress effect

TE (transpiration efficiency) and Wi (intrinsic WUE) were higher in drought conditions, whereas BM (dry biomass), CT (cumulative transpiration), A (net CO_2_ assimilation), and gs (stomatal conductance to water vapor) were lower (Table 1). The effect of drought was not significant for all whole-plant traits measured (see Supplementary File 1): TE, underlying BM and CT were estimated for the whole growing period and for the drought period alone. However, the non-destructive estimation of initial biomass (as used for BM and TE) was more precise (because the plants were smaller and more homogeneous) than estimates of biomass when the desired level of drought stress was reached (as used for BM_d_ and TE_d_). TE, BM and CT, included a period during which the plants were small and not subjected to drought, whereas TE_d_, BM_d_ and CT_d_ covered only the drought period, but may potentially contain more noise due to a less precise initial biomass estimation. This may explain why TE_d_ displayed no significant drought effect, whereas TE, BM_d_, CT and CT_d_ displayed significant drought effect, and BM displayed a tendency at *p*<0.1. A weakly significant interaction was detected for TE_d_ (*p* = 0.046), which was not confirmed by Tukey analysis (Supplementary File 1). We also found no significant effect of drought for BM standardized by leaf area (for the period of drought only and for the whole period), or CT standardized per unit leaf area, but a significant effect was found for standardized CT for the drought period only (Ct_d.LA_). Wi displayed a significant drought effect both *in situ* and in steady state, as did the corresponding stomatal conductance. A_SS_ was also significantly lower under drought conditions, whereas only a tendency was observed for A _in situ_.

**Table 1:**
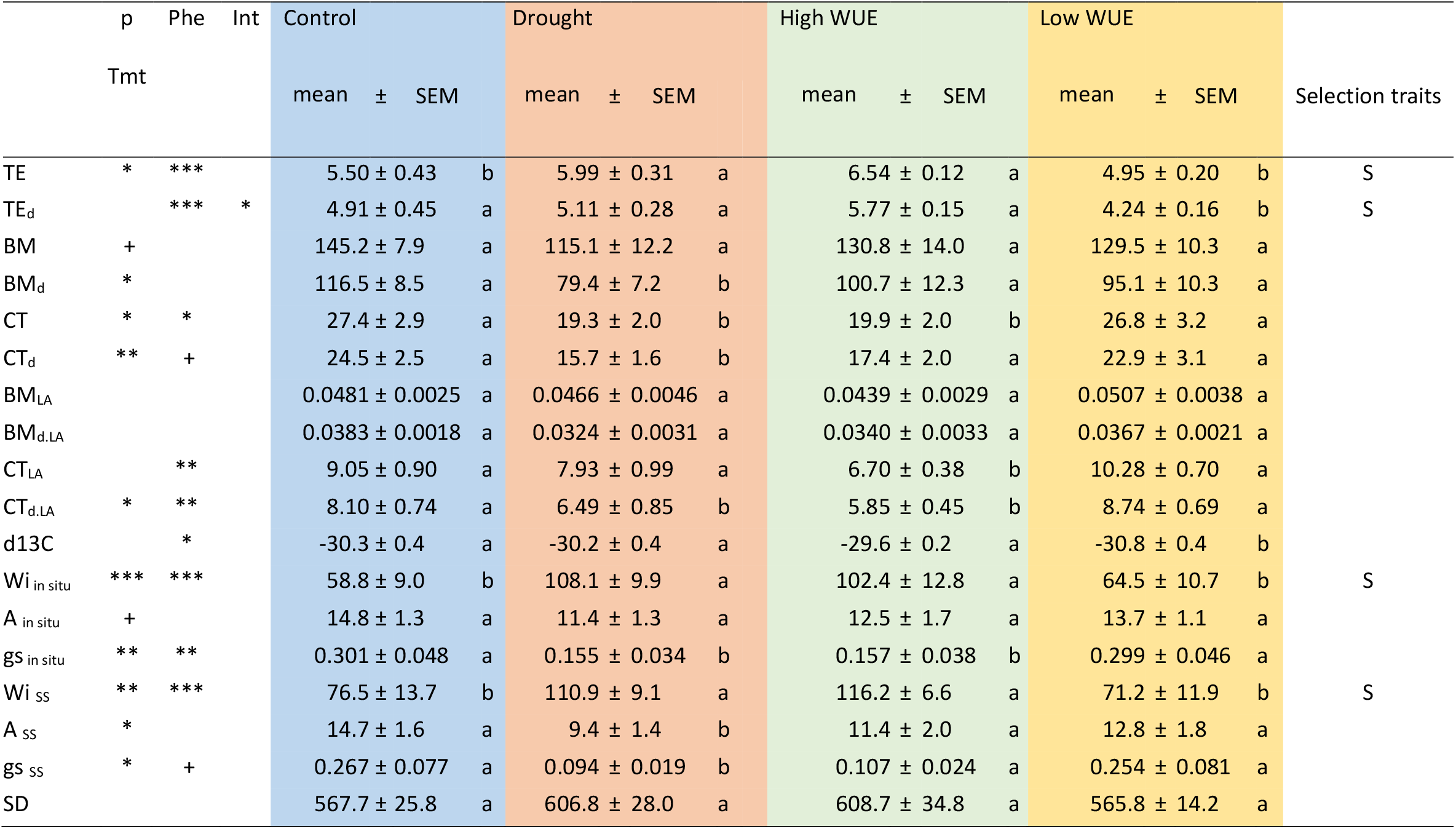
Results of the two-way ANOVA. Means and their standard errors (SEM) for the principal factors tested () and their interaction. Significant differences between groups were identified in Tukey’s highly significant differences test. TE: Transpiration efficiency [g/kg]; BM: dry biomass [g]; CT: cumulative transpiration [kg]; d13C: carbon isotope composition; Wi: intrinsic water use efficiency; A: net CO_2_ assimilation rate; gs: stomatal conductance to water vapor; SD: stomatal density (mm^-1^) indices: d: traits estimated for the drought period only; LA : traits standardized per unit plant leaf area; *in situ*: gas exchange measured under greenhouse conditions; SS: gas exchange performed under steady-state conditions; *p* values: + < 0.1. * < 0.05. ** < 0.01. *** < 0.001; Tmt: watering treatment (control or water deficit); Phe: high- or low-WUE phenotype

#### WUE effect

The selection of sessile oak genotypes based on Wi (Wi_in situ_ and Wi_SS_) and TE (TE and TE_d_) resulted in highly significant differences between the WUE phenotypic groups (Table 1; Figure 1a). Furthermore, the δ ^13^C of bulk leaf material, an independent estimator of integrated Wi, also differed significantly between the high and low WUE phenotypic groups. We investigated the traits underlying TE and Wi. Biomass accumulation (BM, for the whole period or for the period of drought only, and after standardization by leaf area) did not differ significantly between the high and low WUE phenotypic groups. Similarly, no differences were detected for net CO_2_ assimilation rate (A). By contrast, cumulative transpiration (CT) and stomatal conductance (gs) displayed significant differences or strong tendencies in the same direction (CT_d_, gs _ss_), with plants with a high WUE characterized by a lower CT and gs.

### Overview of the cDNA libraries generated

We generated 12 cDNA libraries corresponding to three biological replicates for each of the four groups obtained (2 WUE genotypes x 2 watering regimes).

An overview of the libraries generated is available in Supplementary Table 1. The number of reads ranged from 55 million to 68 million, corresponding to 27 and 34 million paired-sequences, respectively. All the libraries had quality scores above 35. Quality criteria were applied (see methods section), and 26 to 32 million paired reads were finally mapped onto the oak gene models (Supplementary Table 1).

The overall structure of the studied genotypes was investigated by both PCA and hierarchical clustering analysis (Supplementary Figure 1). Unsurprisingly, there was as much variability within as there was between the high and low WUE phenotypic groups. This first observation indicates that only subtle transcriptional changes, if any, drove the variation of WUE between the selected genotypes. This was confirmed by the very small number of DEGs displaying a significant WUE phenotypic effect, consistent with the oligogenic control of WUE (see discussion section). WUE efficiency is driven principally by the density and distribution of stomata [18, 19], and RNAseq analysis was performed on the whole leaf blade, which consists of several cell types. Thus, if the differences in gene expression between genotypes with different WUE values is due to regulation in specific leaf structures, our sampling procedure may have simply diluted the signal, homogenizing the transcription profiles pools of the two groups of six contrasting (high vs. low WUE) genotypes. Despite this cofounding factor, the few DEGs identified were also validated by qPCR, confirming the power of high-throughput sequencing and the robustness of RNA-seq for “finding needles in a haystack”. More striking was the lack of a clear pattern regarding the second structuring factor considered in the factorial design: water availability. This absence of pattern was confirmed by the extremely small number of DEGs associated with watering regime. These findings suggest that the transcriptome of the tissue studied was affected only very mildly by the intensity of the water deficit imposed. The applied, controlled, drought level had been chosen to result in a significant 50% reduction of stomatal conductance (based on gs_in situ_, Table 1), inducing a significant physiological response, but allowing continued growth. After reaching this drought level, it was maintained to over 40 days. Therefor we expected to only detect changes in gene expression with are related to long term, constitutive responses. The different genotypes used in each group might have responded with slightly different strategies, inducing over time more within group variability.

### Identification of differentially expressed genes (DEGs)

We used likelihood ratio tests to identify DEGs displaying significant WUE (high vs. low WUE genotypes), treatment (i.e. watering regime), and interaction effects. This analysis was performed on 15,576 gene models covered by at least five reads in each biological replicate (i.e. 60 reads in total for one gene model). Overall, 92 genes were found to be differentially expressed, with an adjusted *p-*value<0.05. Each DEG presented a unique signature with respect to the main or interaction effects (Supplementary Figure 2).

In total, 12 genes (Gene-set #1, Supplementary File 2) presented a significant “WUE” effect. These genes displayed only quantitative variation. Half were upregulated in the high- or low-WUE genotype. The six genes upregulated in the low WUE phenotypic group were: NAD(P)-linked oxidoreductase super family protein (Qrob_P0056900.2), endoxyloglucan transferase 28 protein (Qrob_P0320680.2), RPL/BEL1-like homeodomain 9 (Qrob_P0256370.2), ALY4 protein (Qrob_P0532680.), nicotinate phosphoribosyltransferase protein (Qrob_P0162910.2) and MAH-like protein (Qrob_P0125520.2).

The six genes upregulated in the high WUE group were: Qrob_P0389480, which is similar to an ankyrin protein, Qrob_P0162930.2, which encodes a protein kinase, Qrob_P0412020.2, which is similar to a DNA-binding protein, a homeodomain-like superfamily protein, Qrob_P0244690.2, which encodes receptor-like protein 15, Qrob_P0489730.2, which is similar to a G-type lectin S-receptor-like serine/threonine-protein kinase, and Qrob_ P0021870.2, which encodes a leucine-rich repeat receptor-like protein kinase family protein.

We also identified 27 genes (Gene-set #2, Supplementary File 3) responding to the watering regime. In total, 17 genes were upregulated in drought conditions, three of which (Qrob_P0268900.2, Qrob_P0104190.2 and Qrob_P0625010.2) were specifically expressed in these conditions. The other 10 genes were upregulated in control conditions.

Finally, 53 genes presented an interaction effect (Gene-set #3, Supplementary File 4) with clear and opposite patterns of expression between the low- and high-WUE phenotypic groups as a function of watering regime. The Kmeans algorithm in Expander software [27] identified two opposite clusters: in cluster #1, 16 genes were upregulated in low-WUE genotypes and downregulated in high-WUE genotypes) with irrigation (i.e. control condition), whereas, in cluster #2, 37 genes displayed the opposite pattern (i.e. upregulated in high-WUE genotypes and downregulated in low-WUE genotypes with irrigation (i.e. control condition)).

Functional annotations for the three gene-sets were retrieved from the TAIR database.

### Analysis of GO term enrichment

We performed GO term enrichment analysis for the three gene-sets described above, corresponding to the main and interaction effects (available in Supplementary Files 2, 3 and 4). The highest degree of molecular function (MF) enrichment was observed for terms: (i) for Gene-set#1 (WUE-responding genes) relating to cytokinin receptor activity, alpha-amylase activity and enone reductase activity, (ii) for Gene-set#2 (drought-responding genes), relating to drug binding, organic cyclic compound binding and heterocyclic compound binding, and (iii) for Gene-set#3 (genes displaying an interaction effect), relating to quercetin 3-O-glucosyltransferase activity, UDP-glucosyltransferase activity and quercetin 7-O-glucosyltransferase activity. For biological process (BP) ontology, we identified terms relating to cellular polysaccharide catabolic processes and ion transport for Gene-set#1, response to external stimulus and cellular processes for Gene-set#2 and defense response and regulation of raffinose metabolic processes for geneset#3.

GO term enrichment analysis was also performed independently for the two contrasting clusters obtained from Gene-set#3. We identified significant enrichment only for cluster#1 (Supplementary File 4) for terms relating to quercetin 3-O-glucosyltransferase activity, UDP-glucosyltransferase activity and glucosyltransferase activity. Enrichment was observed for MF terms relating to the negative regulation of protein catabolic process, and the regulation of vesicle fusion, and for BP terms relating to the regulation of SNARE complex assembly.

### Subnetwork enrichment analysis

We performed subnetwork enrichment analysis independently for the three gene-sets, with Pathway Studio software. Results are also available in Supplementary Files 2, 3 and 4.

For Gene-set#1 (WUE effect), we identified three genes interacting with at least one of the four cellular processes identified (flower development, flowering time, plant development, cell differentiation): a RPL gene (Qrob_P0244690.2) interacting with all four of these processes, an ALY4 gene (Qrob_P0532680.2) and a XTH28 (Qrob_P0320680.2) gene interacting with three of the processes (flower development, plant development and flowering time).

For genes displaying a significant effect of treatment (Gene-set#2), we identified six DEGs known to be involved in five BP. Four of these DEGs (FER (Qrob_P0672440.2), FRK1 (Qrob_P0625010.2), FRO6 (Qrob_P0222260.2) and Akinbeta1 (Qrob_P0268900.2)) were upregulated in the control treatment and regulated three of the five BP identified in this analysis (sugar response, sugar metabolism and chlorophyll content). Two genes (AVP1, (Qrob_P0221580.2) and WOL, (Qrob_P0247500.2)) were overexpressed in the drought treatment and interacted with three BP (turgor, organ formation and chlorophyll content).

We focused in particular on the DEGs displaying a significant WUE-by-treatment interaction effect (Gene-set #3), revealing different molecular strategies for dealing with water constraints in low- and high-WUE genotypes. Subnetwork enrichment analysis was performed for the whole dataset, and separately for clusters #1 and #2. Unfortunately, no significant enrichment was identified for cluster #2. We therefore discuss below only the network obtained for the 53 genes of cluster #1 displaying a significant WUE-by-treatment interaction. We identified nine DEGs (RLK1 (Qrob_P0430150.2), CPR30 (Qrob_P0255490.2), SAG101 (Qrob_P0003420.2), ACA1 (Qrob_P0090430.2), ERH3 (Qrob_P0303640.2), T13D8.29 (Qrob_P0542920.2), VPS41 (Qrob_P0111100.2), UGT74D1 (Qrob_P0574050.2) and CNCG1 (Qrob_P0085080.2)) known to be involved in six BP (regulation of cell shape, plant immunity, cell morphogenesis, photosynthesis, pollen fertility and positive gravitropism). Four of the genes identified (SAG101, 1CA1, CPR30 and ERH3) are involved in regulation of the biological process photosynthesis, and three (RLK1, CPR30 and SAG101) are involved in plant immunity. All the other biological processes identified (pollen fertility, regulation of cell shape, positive gravitropism and cell morphogenesis) are known to interact with two of our DEGs.

### Gene expression – phenotypic trait correlation analysis

A correlation analysis was done between all the ecophysiological traits available in Table 1 and the levels of expression of all genes in the three gene-sets (WUE (#1), Treatment (#2) and Interaction (#3). Given the small number of data points in each regression (*N*=12) and the large number of correlations tested (18 traits and 92 genes, i.e. 1,656 correlations) a stringent type I error of 0.0005 was applied. Eight significant correlations were detected, for six different genes (see supplementary File 5), all with a correlation coefficient above 0.85: Qrob_P0056900.2 was negatively correlated with Wi_SS_ and positively correlated with gs_SS_, Qrob_P0532680.2 was also negatively correlated with Wi_insitu_ and gs_insitu_. Qrob_P0162910.2 and Qrob_P0320680.2 were negatively correlated with TE, whereas Qrob_P0412020.2 was positively correlated with TE_d_. All of these genes belong to geneset#1 (WUE effect). Qrob_P0226850.2 was the only gene from gene-set#2 (Treatment effect) for which a negative correlation was detected with Wi_insitu_ and no significant correlation was detected for geneset#3. All of these genes displayed weaker significant correlations (*p*<0.05, Suppl Table 5) with at least six other traits from Table 1, with the exception of traits related to photosynthesis, for which no significant correlations were detected.

### Real-time quantitative PCR validation

For confirmation of the accuracy of RNA-seq results, we performed reverse transcription-quantitative PCR analyses for 21 genes: six displaying significant WUE effects and 15 displaying WUE-by-treatment effects. Six of the genes displaying significant phenotype-by-treatment interaction effects belonged to Cluster #1 and nine belonged to Cluster #2. We excluded genes displaying either a multiple-band amplification pattern on agarose gels or inconsistent PCR efficiency. We then quantified the expression of the remaining 16 genes by RT-qPCR (5 genes for the phenotype effect, 3 for cluster #1 and 7 for cluster#2, Supplementary Table 2). The patterns of expression of the genes tested by RT-qPCR were similar to those obtained by RNAseq (Supplementary Figure 3), indicating that both the RNA-seq data and the bioinformatics procedure were reliable and that the pattern of DEG expression revealed by RNA-seq could be used for further biological interpretation.

## Discussion

The increase in WUE in drought conditions is due to the stronger effect of drought on stomatal conductance to water vapor than on net CO_2_ assimilation rate. This increase in WUE in leaves can affect transpiration efficiency at whole-plant level [14]. Relationships between leaf and whole-plant level WUE have been documented within various forest tree species [28, 29]. Small changes in functioning at leaf level can, therefore, have major consequences for water use and biomass accumulation. In forest trees, including pedunculate and sessile oaks in particular, strong within-species variability [13, 30] and plasticity [31] related to soil water conditions have been reported for WUE. This suggests that this trait could be used to monitor the capacity of forest genetic resources to maintain sustainable wood production under conditions of water limitation.

The molecular mechanisms involved in regulating WUE are well documented for crops. Abscisic acid seems to play a key role by regulating stomatal movement, thereby increasing WUE [32]. Several studies targeting genes involved in stomatal development and distribution have revealed the biological importance of these processes for WUE regulation and drought tolerance (reviewed in [33]). Other key genes have also been identified. For instance, upregulation of the Hardy gene, encoding an AP2/ERF transcription factor, has been shown to improve WUE by enhancing photosynthesis in rice [21]. More recently, the Erecta gene was identified as a key regulator of WUE in *Arabidopsis*, through the elicitation of changes in leaf epidermal and mesophyll differentiation, with positive effects on growth and biomass accumulation [34]. The elucidation of the molecular mechanisms regulating WUE remains in its infancy for forest trees. In this study, an analysis of differential gene expression enabled us to identify three sets of genes. Below, we focus mostly on the genes of set#1 (i.e. showing a WUE effect, G, supplementary File2) and #3(i.e. displaying a treatment by WUE interaction effect, G*E, supplementary File4), which encode proteins characterizing the molecular strategies of low- and High-WUE genotypes for coping with drought. Such genes may also be involved in the adaptation of forest tree populations to the local variability of water availability.

Our differential gene expression analysis was performed on the whole leaf blade and on different genotypes (i.e. the four WUE by watering treatment combinations are represented by independent genotypes with a specific genetic background). This strategy may account for the relatively small number of DEGs identified. WUE is regulated principally by stomatal aperture and CO_2_ carboxylation, in a process involving a relative small number of genes [17] with respect to the total transcriptome expressed in a plant. One previous study [15] reported an oligogenic genetic determinism of WUE in a pedunculate oak family, supporting the hypothesis that trait variation is influenced by a small number of genes, each with a large effect, consistent with our observations for oak (see the high fold-change ratios below). The subnetwork enrichment analysis identified key genes of potential importance for sessile oak WUE (see below), demonstrating that the RNaseq approach is a method of choice for accelerating gene discovery for fine ecophysiological traits, as reported in many crops (reviewed in [35]), even in the presence of cofounding and homogenizing effects. Our RNAseq data were also validated by RT-qPCR.

### (i) Genes regulated by soil water deficiency

In total, 27 genes in Gene-set#2 were found to correspond to the effect of soil water deficit, and the subnetwork enrichment analysis identified BPs (turgor, organ formation, chlorophyll content and sugar metabolism) known to be regulated by drought stress [36]. The overexpression of two genes (WOL, FC=0.22 and AVP1, FC=0.28) regulating turgor in plants is consistent with several studies that have already reported that cell turgor regulation during water depletion is essential to cope with stressful conditions [19]. We also observed a decrease in transcript accumulation for four genes involved in sugar metabolism, sugar response and chlorophyll content, probably due to the decrease in primary metabolism classically observed during the abiotic stress response [37]. We found a highly significant correlation between Qrob_P0226850.2 (FC=2.24), which encodes an abscisic acid-responsive gene (AT1G02130), and intrinsic water use efficiency, highlighting the coordination of the drought response by ABA at leaf level.

### (ii) Genes regulated in high and low WUE Genotypes

Of considerable relevance to our research question, the second set included 12 genes constitutively expressed at higher levels in either low- or high-WUE genotypes (Gene-set#1, Supplementary File 2) regardless of the watering regime, providing clues to the molecular mechanisms underlying high WUE in sessile oak.

Also, in total, six genes were sinificantly correlated with TE, Wi or stomatal conductance (Supplementary file 5), which were all also detected for constitutive expression differences (Supplementary file 2). These genes were not correlated with assimilation rate, suggesting that the genetic diversity of WUE observed in these sessile oak genotypes is driven principally by variation in stomatal opening. This is coherent with what has already been shown in a Q. robur full-sib family [14,16]. The correlation analysis also showed that the expression of Qrob_P0056900.2, which is similar to an NAD(P)-linked oxidoreductase superfamily protein (AT1G59960), was inversely correlated with WUE and positively correlated with gm SS.

Six genes upregulated in high WUE individuals were considered of particular interest: (i) the receptor-like protein 15 gene (RPL15, Qrob_P0244690.2,) which presented a 4.9-fold change in expression. RPL genes encode cell-surface receptors involved primarily in defense against pathogens. Some RPL genes are also known to regulate stomatal density and distribution by disrupting their patterning. For example, the too many mouth (TMM) gene is highly similar to RLP16 [38]. Other authors have identified TMM as a key gene for improving the WUE of crops through molecular genetics approaches [17]; (ii) we also identified a leucine-rich repeat receptor-like protein kinase gene (Qrob_P0021870.2, validated by RT-qPCR) displaying a 5.2-fold change in expression. Leucine-rich repeat receptor-like protein kinases are receptor kinases located on the surface of the plant cell and involved in the perception of signals from the environment. A study in *Arabidopsis thaliana* by [39] reported a key role for this gene in both regulating several aspects of growth and the response to abiotic stress; (iii) we identified a G-type lectin S-receptor-like serine/threonine protein kinase gene (Qrob_P0489730.2, validated by RT-qPCR) with a 4.8-fold change in expression. The overexpression of this gene in *Arabidopsis* [40] enhances salinity and drought stress tolerance; (iv) a homeodomain-like superfamily protein (Qrob_P0412020.2)gene, encoding a DNA-binding protein, was also identified, with a 2.02-fold change in expression, for which expression was significantly correlated with TEd (and, to a lesser extent, other estimators of WUE and whole-plant transpiration). In *Quercus Lobata*, it has been suggested that such genes may underlie the differences in WUE between oak populations [41]; and (v) finally, we identified an ankyrin protein gene (Qrob_P0389480.2) with a 4.7-fold change in expression. Ankyrin repeat proteins play essential roles in cell growth, development and the response to hormones and environmental stresses [42]. It has recently been reported [43] that some ankyrin genes confer resistance to drought and salinity in *Arabidopsis*, probably by regulating water metabolism. Five of the six genes upregulated in low-WUE genotypes (or down-regulated in high-WUE genotypes) were considered of considerable interest based on published findings or the results of our subnetwork analysis:

(i) an MAH-like protein gene (−0.83-fold change in expression, Qrob_P0125520) potentially involved in wax and cutin biosynthesis [44]. It has been suggested that changes in cutin thickness may be associated with WUE variation in pine, through effects on transpiration rate [45]. However, no clear relationship is available in the literature on the influence of cuticle thickness on WUE. (ii) the NAPTR1 gene (−0.32-fold change in expression, Qrob_P0162910), which encodes a nicotimate phosphoribosyltransferase known to play a key role in both the biosynthesis and homeostasis of NAD [46]. NAD homeostasis has recently been shown to be a key component in stomatal movement. The authors also showed, with transgenic plants, that mutations of this gene led to the stomata partially losing their ability to close, resulting in higher drought stress sensitivity [47]. NAPTR1 expression was inversely correlated with TE (and, to a lesser extent, with other estimators of WUE, transpiration and stomatal conductance), suggesting a major role in stomatal opening, driving differences in WUE. The three last genes were associated with the functional network (Supplementary File 2): (iii)ALY (−0.26-fold change in expression, Qrob_P0532680) genes encode key proteins involved in the transport of mRNA from the nucleus to the cytosol and are essential for correct plant growth and development [48]. It has also been shown [49] that the ALY4 gene is involved in stomatal closure in *Arabidopsis*. We also observed two highly significant and strong correlations between ALY4 expression and Wi_insitu_ (inverse correlation) and to gs_intisu_ (positive correlation), suggesting that higher levels of expression for this gene resulted in greater stomatal opening in low-WUE genotypes, (iv) the XTH28 (−0.71-fold change in expression, Qrob_P0320680.2) gene encodes a xyloglucan endotransglycosylase (XTH) protein. In plants, the members of the XTH multigene family are involved in the metabolism of xyloglucan, an important cell-wall compound [50]. In *Solanum lycopersicum*, a role for the product of the XTH3 gene in regulating cell wall remodeling in guard cells, and, potentially, in regulating transpiration rates during abiotic stress has been reported [51], suggesting a potential role of this gene in WUE. A significant correlation was observed between the expression of this gene and the whole plant transpiration efficiency (TE), and to a lesser level with 10 other traits, all related to stomatal conductance and cumulative water consumption (positive correlations), as well as whole plant (Te_d_) and integrated (δ ^13^C) WUE (negative correlations). This analysis suggests that a stronger expression of this gene will decrease WUE by increaseing stomatal conductance and thereby the cumulative water consumption. (v) we identified an RPL gene encoding a Bel 1-like homeodomain (−2.14-fold change in expression, Qrob_P0256370). In rice, Bel1 has been implicated in the regulation both of organ size and water loss, through the regulation of stomatal density. In studies of mutations of Bel1 genes [52], a key role was identified for the product of the RPL gene in cell proliferation, expansion and response to environmental clues. A higher stomatal density, leading to senescence associated with underdeveloped chloroplasts and low levels of photosynthesis, was observed in this mutant, consistent with a greater sensitivity to drought stress.

The identification of these DEGs highlights a series of genes for which up- or downregulation is associated with high WUE in sessile oaks regardless of the watering regime. An overview of the genes identified is available in Figure 2, with the corresponding BPs, which include stomatal density and distribution, cuticle biosynthesis, regulation of transpiration or stomatal movement. This study thus validates current scientific knowledge regarding the importance of water loss regulation in genotypes with a high water-use efficiency, including those of sessile oak.

**Figure 2:**
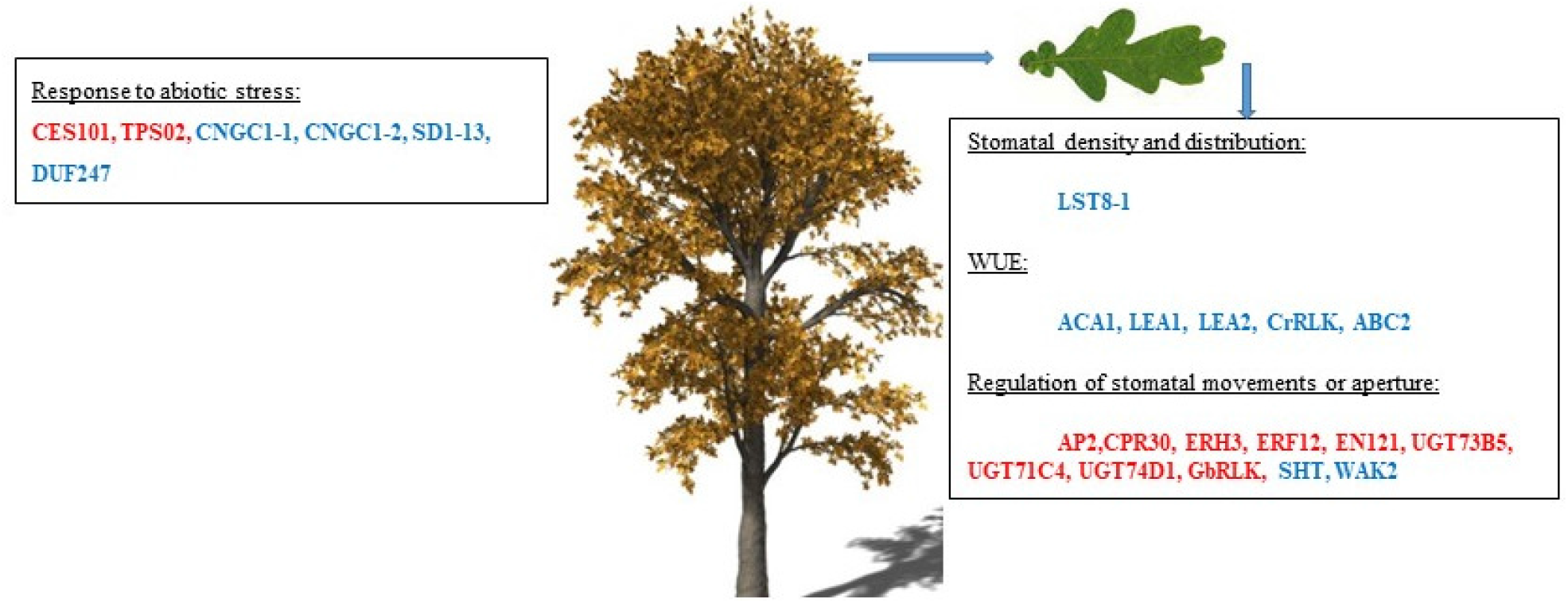
Illustration of the main biological processes and associated genes regulated by WUE in sessile oaks.

### (iii) Gene regulation triggering high WUE under drought conditions: toward adaptive molecular mechanisms combining drought tolerance and WUE

The third set of genes, displaying GxE (i.e. corresponding to genes for which plasticity in gene expression was dependent on the WUE of genotype studied), is of even greater interest in the context of this study, as these genes represent key candidate genes of importance in the adaptation of oak to water scarcity.

Two clusters were identified by the Kmeans approach. Cluster #1 contained 16 genes downregulated in the low-WUE genotypes and upregulated in high-WUE genotype in drier conditions. Cluster #2 comprised 37 genes with the opposite pattern of expression (i.e. up- and downregulated in the low- and high-WUE genotypes, respectively, in drier conditions). These two clusters of genes correspond to molecular mechanisms enabling oak trees to maintain a high WUE in conditions of water deficit. They may, therefore, correspond to key genes enabling plants both to regulate their transpiration rate and to maintain their carbon-fixation capacity in a drier environment. The identification of genes associated with a high WUE in a drier environment is of particular interest for the management of forest plantations in the context of global warming. This gene pool may, therefore, correspond to genes of importance for the adaptation of forests to a drier environment, therefore worth for conserving their genetic diversity. A description of the biological processes and associated genes identified is available in Figure 3.

**Figure 3:**
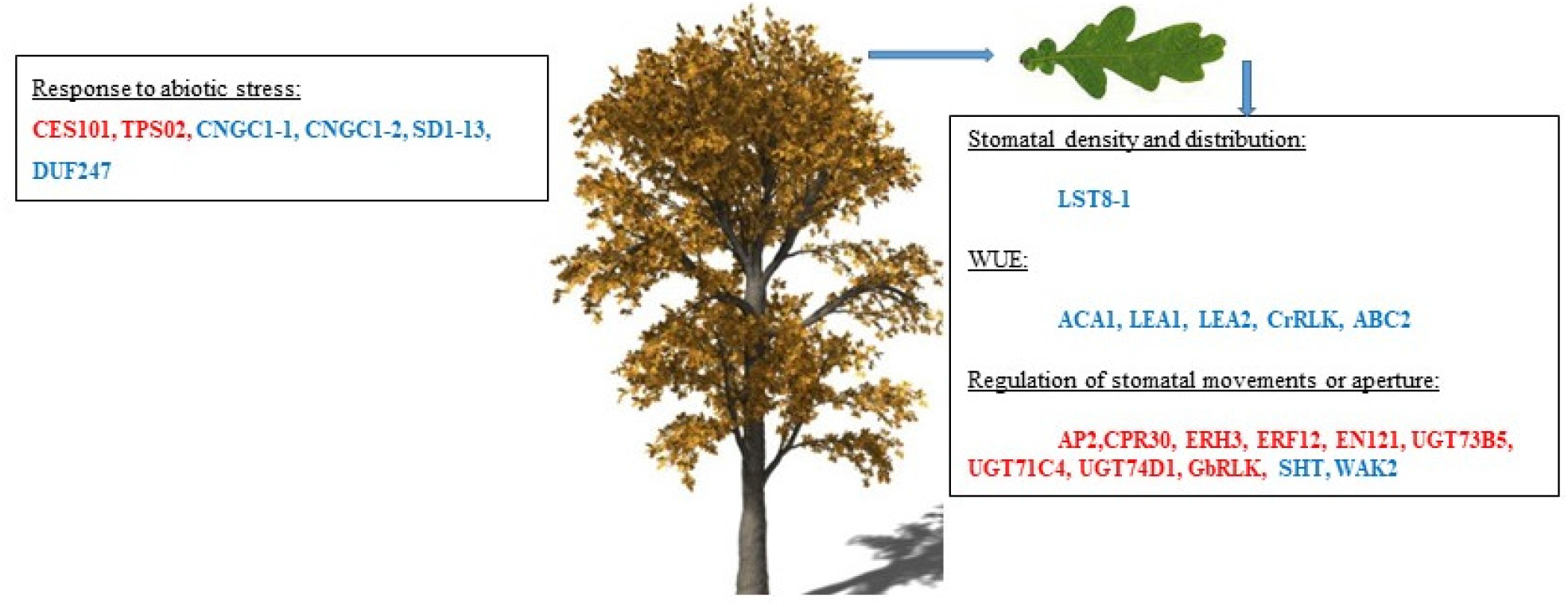
Illustration of the main biological processes and associated genes triggering high WUE under drought stress in sessile oak.

In cluster #1, we identified two genes validated by RT-qPCR (CES101, Qrob_P0587340.2 and TPS02, Qrob_P0198710.2) and known to be regulated by drought stress. The first, CES101, is highly similar to a lectin S-receptor-like serine/threonine protein kinase, the expression of which is modulated during salt stress in *Arabidopsis thaliana* [53]. The second, TPS 02, encodes a terpene synthase. Monoterpenes are important aromatic molecules widespread in nature. The levels of monoterpene and terpene synthases increase massively in plants exposed to drought stress [54]. The upregulation of a TPS gene in high-WUE genotypes may result in better drought tolerance. We identified eight other genes (AP2 (Qrob_P0748930.2), CPR30 (Qrob_P0255490.2), ERH3 (Qrob_P0303640.2), ERF12 (Qrob_P0492110.2), EN121 (Qrob_P0638250.2), and three UGT genes (UGT73B5 (Qrob_P0430640.2), UGT71C4 (Qrob_P0125190.2), UGT 74D1(Qrob_P0574050.2)) known to be involved in the regulation of stomatal movement or stomatal aperture by regulating IAA levels in guard cells [55]. For example, AP2 is similar to aspartic protease 2, which is specifically expressed in guard cells. In *Arabidopsis thaliana*, overexpression of the AP2 gene resulted in a smaller mean stomatal aperture, suggesting a possible role for the AP2 gene in the fine regulation of transpiration rate [56] and, indirectly, in WUE. EN121, another of the genes identified, encodes a basic helix-loop-helix (bHLH) DNA-binding protein. In wheat, bHLH proteins are transcription factors known to be regulated by both ABA and drought stress. A transgenic line overexpressing the bHLH gene was found to display early stomatal closure in response to drought stress, enabling the plant to regulate its transpiration more effectively [57]. We identified an ERF12 gene displaying similarity to ethylene-responsive factor 12 gene. A transgenic line of cotton overexpressing the ERF38 gene displayed both a larger guard cell stomatal aperture and a lack of sensitivity to ABA in terms of stomatal closure under drought stress [58], suggesting a key role of ERF in regulating transpiration rate. The last two genes (CPR30 and ERH3) were identified by functional network analysis (Supplementary File 3). The ERH3 gene is similar to a katanin P60 subunit protein. In *Arabidopsis thaliana*, an ERH3 loss-of-function mutant had small stomata, suggesting a possible role for the katanin protein in both cell differentiation and stomatal development [59]. We hypothesized that the differential expression of genes involved in stomatal size would affect WUE by regulating both transpiration and assimilation rate. However, no significant differences in stomatal size were observed in our study. Finally, we identified a CP30 gene highly similar to an F box protein as differentially expressed. The CPR 30 gene was also found to be upregulated in response to drought stress in another European white oak, *Quercus pubescens* [60]. CPR30 is known to interact with ASK proteins, which promote stomatal opening, suggesting a possible functional role of CPR30 in the regulation of WUE [61], by increasing the transpiration rate under drought stress. Finally, we identified three UGT genes (UGT73B5, UGT71C4 and UGT74D1). UGT74D1 is a key gene involved in auxin glycosylation, a key mechanism regulating the levels of auxin in cells [55]. Indole-3-acetic acid (IAA) is a key auxin hormone involved in the regulation of several biological processes. It has been reported to stimulate stomatal opening [62]. The regulation of IAA levels by genes in cluster #1 may explain the lower WUE, through the modulation of stomatal movements. However, further studies would be required to confirm this, as no IAA determinations were performed in this study.

Fourteen of the 37 genes in cluster #2 were considered of potential interest. Indeed, all these genes are known to be involved in drought stress tolerance (4 genes) or in the regulation of stomatal movements and/or WUE (10 genes). We first identified genes known to improve resistance to drought stress: two CNGC1 genes (Qrob_P0085080.2 validated by RT-qPCR and Qrob_P0274450.2) and an SD1-13 (Qrob_P0394960.2, validated by qPCR) gene known to upregulated during drought stress in *Arabidopsis thalian*a [63, 64]. We also identified a DUF 247 gene (Qrob_P0059340.2, validated by RT-qPCR) known to be differentially regulated in response to drought stress *Quercus lobata* [41]. The other 10 genes have been implicated in WUE and/or in the regulation of stomatal movement. We identified an ACA1 gene (Qrob_P0090430.2) encoding a carbonic anhydrase. ACA genes encode key enzymes involved in the transfer of CO_2_ to the catalytic site of ribulose 1.5 biphosphate carboxylase/oxygenase in the cell, which may act as important regulators of WUE in plants [65]. We identified two late embryogenesis abundant (LEA) proteins (Qrob_P0273440.2 and Qrob_P0714290.2). Barley transgenic lines with LEA mutations perform better in terms of both biomass productivity and WUE under drought stress [66]. We also identified two genes encoding RLK proteins (Qrob_P0430150.2, CrRLK and Qrob_P0210860.2, GbRLK, validated by RT-qPCR). In rice, CrRLK proteins are known to be involved in the regulation of the circadian clock and, potentially, in WUE [67]. GbRLK proteins are key cell-surface receptors that may regulate stomatal movements in response to environmental cues in cotton [68]. Indeed, several author reported that loss of function mutant for some GbRLK gene were defective for stomatal closure [69]We identified an SHT protein in this cluster that was upregulated (Qrob_P0064930.2, validated by RT-qPCR). SHT proteins are involved in the polyamine biosynthesis pathway. Several authors have reported that polyamines are key components of the plant response to environmental variations. Indeed, they may be involved in stomatal movement through regulation of the voltage-dependent inward K^+^ channel in the plasma membrane of the guard cell [70]. We identified an ABC2 gene (Qrob_P0453640.2, validated by RT-qPCR) encoding an ATP-binding cassette. In *Arabidopsis*, loss-of-function mutations affected WUE under drought stress, suggesting a key role for this gene in water uptake and loss [17, 71]. We also identified a WAK2 gene encoding a serine/threonine kinase (Qrob_P0186810.2). In *Arabidopsis thaliana*, loss-of-function mutations of this serine/threonine kinase gene strongly decreased WUE, suggesting a possible role for this gene in the regulation of stomatal movement [72]. Although the role of this gene under drought conditions is not very clear for high WUE genotype, its upregulation in control conditions may explain the Higher WUE observed. Finally, we identified two LST8-1 genes (Qrob_P0168200.2 and Qrob_P0168190.2). LST8-1 is a partner of the target of rampamycin kinase involved in several biological processes in plants. The LST8-1 gene was recently reported to be strongly expressed in *Arabidopsis* guard cells, suggesting a possible in stomatal regulation [73].

## Conclusion

We applied an RNAseq approach to 12 independent sessile oak genotypes differing in terms of WUE, to decipher the molecular mechanisms underlying WUE variation, and to determine which of these mechanisms contribute to the maintenance of high efficiency under drought stress.

We constructed a schematic model summarizing the main findings (Figures 2 and 3, based on the data obtained in this study. According to this model, genotypes with a high WUE display regulation of the constitutive expression of 12 genes (6 upregulated and 6 downregulated). Under drought stress, they maintain their growth capacity upregulating 16 genes and downregulating 37 genes.

We first identified genes differentially regulated between individuals with contrasting WUE (low *vs*. high WUE, Gene-set#1). Low-WUE genotypes displayed a regulation of genes involved in cuticle remodeling or the rapidity of stomatal closure in response to drought stress, whereas high-WUE genotypes were characterized by a regulation of genes involved in stomatal density and distribution, or genes encoding surface receptors potentially involved in the regulation of transpiration. These genes may contribute to fine regulation of the tree’s transpiration rate, improving WUE in these genotypes.

We also identified genes displaying a significant WUE-by-treatment interaction, highlighting different molecular strategies between low- and high-WUE genotypes for coping with water shortage. The expression profile of low-WUE genotypes highlighted an important role for genes involved in early stomatal closure or in increasing transpiration rate, whereas that obtained for high-WUE genotypes highlighted an important role for genes involved in the carboxylation of CO_2_and genes involved in the regulation of stomatal movements or in water uptake and loss under drought stress. These genes may enable sessile oak seedlings to keep their stomata open, thereby maintaining their transpiration rates during drought stress and resulting in a higher WUE.

This comprehensive investigation constitutes a first step toward understanding the molecular basis and adaptive value of WUE. Additional insights are emerging from ongoing investigations focusing on the association between WUE and gene polymorphisms in a pedigree population and association panels of unrelated sessile oak genotypes.

## Methods

### Plant material and experimental design

The plants analyzed here were part of a larger experiment, carried out at INRAE (Champenoux, France, 48°45’8’’N, 6°20’28’’E, elevation: 259 m asl) on 60 *Q. petraea* seedlings originating from six different French provenances. The seeds were selected from stands growing in conditions ranging from dry to moderately humid, to capture a maximum of the within-species variation. The acorns were collected in autumn 2015 and sown during spring 2016 in 6 L pots filled with a 5/3/2 (V/V/V) mixture of sand, peat and silty-clay forest soil. This soil mixture was tested to a field capacity (FC) of 33% soil water content (SWC). The plants were grown in a greenhouse equipped with a robotic system for the automatic weighing and watering of plants [74]. All the plants were initially subjected to the same non-limiting growth conditions: natural light, with fertilization and irrigation at 85% of relative extractable water (REW, with field capacity at 33% and a wilting point at 4% soil volumetric humidity). The volumetric soil water content was measured by regular time domain reflectrometry (TRIME-TDR; IMKO GmbH, Ettlingen, DE), with one to two measurements per week.

### Measurement of soil water status and establishment of a water deficit

Plants were randomly assigned to the control and drought groups at day 193, with the REW decreased to 31% REW for the water stress group (reached at day 221). The final harvest took place during day 266. For more details see [75].

### Monitoring of leaf gas exchange

Regular *in situ* gas exchange monitoring was performed with a portable photosynthesis system (Li-Cor 6200; Li-Cor, Lincoln, NE, USA). Net CO_2_ assimilation rate (A_m_), and stomatal conductance to water vapor (g_m_) were measured four times during the drought period, on the same third-flush leaf. Leaf intrinsic water use efficiency (Wi) was estimated as A/g, and an overall mean of all measurements during the drought period was then calculated (Wim).

Leaf gas exchange was also measured in a more standardized way with a Li-COR Cor 6400XT (LiCor, Lincoln, NE, USA) equipped with a 2 cm^2^ leaf chamber. All measurements were performed between 10:00 and 19:00 h (Central European summertime), with the following conditions in the leaf chamber: CO_2_ concentration of 400 µmol mol^-1^, temperature of 25°C, ∼70% humidity, and a PAR of 1200 µmol m^-2^ s^-1^. These measurements were performed three times during the drought period for each plant. Wi was calculated as A/g and an overall mean was then calculated (Wis).

#### Final harvest

At the end of the experiment on the 266^th^ day of the year, all the plants were harvested. Plants were oven-dried at 60 °C until they reached a constant dry mass. Leaves were weighed separately to estimate the total leaf area (LA) according to the following allometry established in a previous study [75]: LS= 125.67*LBm+574.59 (rR^2^ = 0.82, *n* = 30)

The leaves used for standardized gas exchange were harvested and cut in half. One half of the leaf was used for stable carbon isotope analysis, and the other half was used for stomatal density measurements. The leaves for bulk leaf carbon isotope composition (δ ^13^C) were ground in a ball mill and analyzed with an elemental analyzer (vario ISOTOPE cube, Elementar, Hanau, Germany) coupled to an isotope ratio mass spectrometer (IsoPrime 100, Isoprime Ltd, Cheadle, UK). We calculated δ^13^C according to the international standard (Vienna Pee Dee Belemnite, VPDB).

#### Measurement of stomatal density

Oak trees have stomata only on the abaxial surface of the leaves. We collected 1cm^2^ segments of leaf and took nail varnish imprints of the abaxial surface with an adhesive film, which was then applied to a microscope slide for analysis. ImageJ2 software was used to count the stomata on the images obtained, excluding stomata overlapping the margins of the image.

#### Estimation of transpiration efficiency

For the estimation of TE, the total water consumption (TWC) of the plants was measured by adding the amount of water from all the watering cycles of each plant over a four months period from mid-May to the final harvest in September, corrected for the water evaporation from pots without plants (H2Oall). This calculation was also performed for the drought period only (H2Od). Plant water consumption is also expressed on a per unit leaf area basis (H2OallLA and H2OdLA), based on LA at the end of the experiment. Initial dry biomass, in mid-May was estimated from height measurements, according to an allometric relationship [75]. We also estimated dry biomass at the start of the drought period. The increase in biomass was calculated for the whole period (BMall) and for the drought period only (Bmd). TE efficiency was calculated by dividing accumulated biomass by water consumption for the whole period (TEall) and for the drought period only (Ted).

### Selection of plants for RNAseq

At the final harvest, just before the plants were oven-dried, third-flush leaves close to those used for gas exchange were harvested from all 60 plants and frozen in liquid nitrogen for RNA extraction. We selected 12 plants, on which we measured Wi _insitu_, Wi _ss_, TE, and TE_d_. Individual trees were chosen so as to constitute high- and low-WUE phenotypic groups consistent for all four measurements of WUE. First, for each trait, the upper and lower quantiles were chosen (separately for control and drought plants). The trait values were transformed into ranks and the mean rank was calculated. The six plants with the most extreme ranks in each group were selected. Finally, three plants per group were chosen, by visual selection, to constitute groups homogeneous for all traits.

### Statistical analysis of physiological traits

All statistical analyses were performed with R 3.4.3 (R Core Team (2015). The significance of differences between treatments were analyzed by one-way analysis of variance (*t*-test, *n* = 60). Differences were considered significant if *P* < 0.05.

A Principal components Analysis (PCA) was performed for the four traits used for selection (TE, TE_d_, Wi_in situ_,Wi_SS_) and also for all traits that were measured (N=18).

### RNA extraction, library construction and sequencing

We first carefully removed all leaf veins, to focus the analysis exclusively on the leaf blade. RNA was then extracted as previously described [76]. Total RNA was recovered from the 12 selected genotypes with a CTAB-based extraction buffer. Residual genomic DNA was then removed from each sample with 10 U of RNAse-free DNaseI (Promega, Madison, WI, USA) according to the manufacturer’s instructions. After DNAse digestion, the RNA was purified with the cleanup protocol from the RNeasy plant minikit (Qiagen, Valencia, CA, USA). RNA quantity and quality was assessed for each sample by both optical density measurements and electrophoresis on an Agilent 2100 Bioanalyzer (Agilent Technologies, Inc., Santa Clara, CA, USA). RNA integrity number (RIN) values ranged from 7 to 8.5 for the 12 samples. We then generated cDNA libraries from 2 µg of RNA according to the Illumina protocol (TruSeq Stranded mRNA Sample Prep Kit). We sequenced each library with 150 base reads, in a paired-end flow cell, on an Illumina HiSeq2000 (Illumina, San Diego, CA, USA). Over 50 million useable reads were generated for each library (Supplementary Table 1). The sequences were produced by Genewiz (Leipzig, Germany).

### Trimming, mapping and identification of differentially expressed genes (DEGs)

Bioinformatic analyses were performed in a Galaxy web environment according to a published procedure [77]. The quality of each batch of sequences was first assessed with FAstQC, and low-quality reads (i.e. quality value<20) were removed. High-quality reads were then mapped onto the 25k pedunculate oak gene models [78] with BWA V0.6.1 [79]. We finally selected only gene models covered by at least 60 reads (i.e. at least 5 reads per sample) for differential gene expression analysis. DEGs were identified with DESeq2 software [80], with a *p*-value<0.05 after adjustment for multiple testing with a false discovery rate (FDR) of 5%. The effects of WUE (i.e. low vs. high WUE), treatment (i.e. control vs. drought) and their interaction were assessed in likelihood ratio tests implemented in the DESeq2 package. The WUE and treatment effects were investigated by comparing a model without interaction (M1) with two reduced models for the treatment (M2) and WUE (M3) effects. For estimation of the interaction effect, we compared M4 to M1.

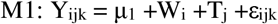

where W_i_ is the WUE effect (i= “high WUE” vs. “Low WUE”), T_j_ is the treatment effect (j= “Control” vs. “drought”).

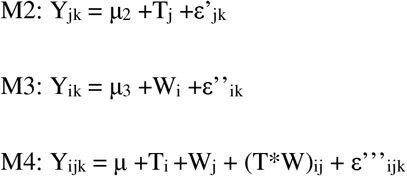

Only three of the seven theoretical categories of genes displaying a single or combined effect (WUE, treatment or WUE x treatment interaction) included DEGs. Their annotations were recovered from the corresponding oak gene models [78].

#### Gene set and subnetwork enrichment analysis

GO term enrichment analysis was performed for the DEGs within each of the three categories, with topGO in R [81]. We corrected the *P*-value for false discovery rate and considered ontology terms with a corrected *p*-value below 0.05 to be significantly enriched. Results were displayed for the first 10 ontologies only.

We then performed pathway enrichment analysis with Pathway Studio software (Pathway Studio®, Elsevier 2017), as previously described [77]. We considered a *P*-value<0.05 to indicate significant enrichment in a biological process (BP) or a molecular function (MF) in our DEG datasets. Gene networks were then generated with the gene sets corresponding to the main and interaction effects.

#### RT-qPCR validation

We quantified the expression of 21 DEGs. All the primer pairs used for the qPCR were designed with Primer3 software [82]. All the genes analyzed are listed, together with their associated effects, in Supplementary Table 2. For each RT-qPCR reaction, we normalized the fluorescence data against two control genes (Qrob_P0530610 and Qrob_P0426000) selected from the non-DEG and characterized by highly repeatable levels of expression (i.e. similar numbers of reads across the different samples analyzed). qPCR was performed on a LightCycler480 Real-Time PCR detection system (Roche Applied Science, Penzberg, Germany) with standard PCR parameters, as previously described [77]. The fluorescence data obtained were analyzed with StatQPCR software, using methods derived from the Genorm and qBASE programs. StatQPCR is available from the following URL: http://satqpcr.sophia.inra.fr/cgi/tool.cgi.

## Supporting information

Overview of the cDNA libraries constructed in this study.

PCA analysis and clustering analysis of the data generated in this study.

Venn diagram showing the overlap between the three effects analyzed in this study.

qPCR validation of the candidate genes selected in this study.

List of the primer pairs used for qPCR analysis.

## Declarations

Availability of data and materials: The datasets generated and/or analyzed in this study are available from the SRA-NCBI repository under accession number PRJNA763825. All the scripts used for bioinformatics analysis and the identification of differentially expressed genes are available from the corresponding author upon request.

## Authors’ contributions

GLP, TG and OB designed the study. GLP and CP wrote the manuscript with OB. GLP performed bioinformatics analysis. GLP was involved in the RT-qPCR experiments. OB and TG were responsible for running the experiment and the ecophysiological measurements and their statistical analysis. All the authors have read and approved the manuscript.

## Acknowledgments

We thank the Genotoul bioinformatics facility in Toulouse for providing computing resources. qPCR analysis was performed at the sequencing and genotyping facility of Bordeaux, supported by Conseil Régional d’Aquitaine grants nos. 20030304002FA and 20040305003FA, European Union FEDER grant no. 2003227 and Investissements d’Avenir ANR-10-EQPX-16-01. We also thank Cyril Buré for help with the preparation and the running of the experiment and the management of the robotic irrigation system.

## Supporting Information

**Supplementary Tables 1**: Overview of the cDNA libraries constructed in this study.

**Supplementary Table 2**: List of the primer pairs used for qPCR analysis.

**Supplementary Figures 1**: PCA analysis and clustering analysis of the data generated in this study.

**Supplementary Figure 2** Venn diagram showing the overlap between the three effects analyzed in this study.

**Supplementary Figure 3**: qPCR validation of the candidate genes selected in this study.

**Supplementary Files 1, 2, 3, 4, 5are available online INRAE dataverse: G. Le Provost, T. Gerardin, C Plomion, O. Brendel Molecular plasticity to soil water deficit differs between sessile oak (Quercus Petraea (Matt.) Liebl.) high- and low-water use efficiency genotypes**, https://doi.org/10.15454/J1VQDO **Portail Data INRAE**,**V3.0**. **These files includes normalized values for RNAseq data, Fold change Ratio and Gene set enrichment analysis as well as ecophysiological measurement and correlation analysis (File 4 and 5)**

