## Supplementary material for "Molecular plasticity to soil water deficit differs between sessile oak (*Quercus Petraea* (Matt.) Liebl.) high- and low-water use efficiency genotypes": Overview of the cDNA libraries constructed in this study.

Supplementary Table 1: Overview of the cDNA libraries generated in this study. Abbreviations correspond to: HighWUE, High Water Use Efficiency phenotype, LowWUE, Low Water Use Efficiency phenotype.

| **Library** | **Phenotype** | **Treatment** | **species** | **Biological replicate** | **Number of read** | **Number of read mapped** | **SRA-NCBI study Accesion** |
| --- | --- | --- | --- | --- | --- | --- | --- |
| 2 | **HighWUE** | **Control** | ***Quercus Petraea* Matt L.** | **Bio. Rep. 1** | 63,5462,52 | 38,660,374 | PRJNA763825 |
| 29 | **HighWUE** | **Control** | ***Quercus Petraea* Matt L.** | **Bio. Rep. 2** | 64,626,871 | 41,377,008 |  |
| 30 | **HighWUE** | **Control** | ***Quercus Petraea* Matt L.** | **Bio. Rep. 1** | 67,6406,84 | 42,654,916 |  |
| 5 | **HighWUE** | **Drought** | ***Quercus Petraea* Matt L.** | **Bio. Rep. 2** | 62,452,217 | 39,167,906 |  |
| 25 | **HighWUE** | **Drought** | ***Quercus Petraea* Matt L.** | **Bio. Rep. 1** | 68,410,305 | 42,501,006 |  |
| 59 | **HighWUE** | **Drought** | ***Quercus Petraea* Matt L.** | **Bio. Rep. 2** | 57,981,714 | 36,478,742 |  |
| 20 | **LowWUE** | **Control** | ***Quercus Petraea* Matt L.** | **Bio. Rep. 1** | 56,177,197 | 46,196,958 |  |
| 37 | **LowWUE** | **Control** | ***Quercus Petraea* Matt L.** | **Bio. Rep. 2** | 60,686,048 | 38,226,472 |  |
| 47 | **LowWUE** | **Control** | ***Quercus Petraea* Matt L.** | **Bio. Rep. 1** | 63,499,984 | 41,207,444 |  |
| 16 | **LowWUE** | **Drought** | ***Quercus Petraea* Matt L.** | **Bio. Rep. 2** | 55,808,270 | 36,617,750 |  |
| 19 | **LowWUE** | **Drought** | ***Quercus Petraea* Matt L.** | **Bio. Rep. 1** | 62,567,776 | 40,349,226 |  |
| 56 | **LowWUE** | **Drought** | ***Quercus Petraea* Matt L.** | **Bio. Rep. 2** | 59,893,313 | 38,863,748 |  |
