## Supplementary material for "Molecular plasticity to soil water deficit differs between sessile oak (*Quercus Petraea* (Matt.) Liebl.) high- and low-water use efficiency genotypes": PCA analysis and clustering analysis of the data generated in this study.

Supplementary Figure 1: PCA analysis (panel A) and clustering analysis (using the “complete” method of the expender software, panel B) among the 12 different libraries.

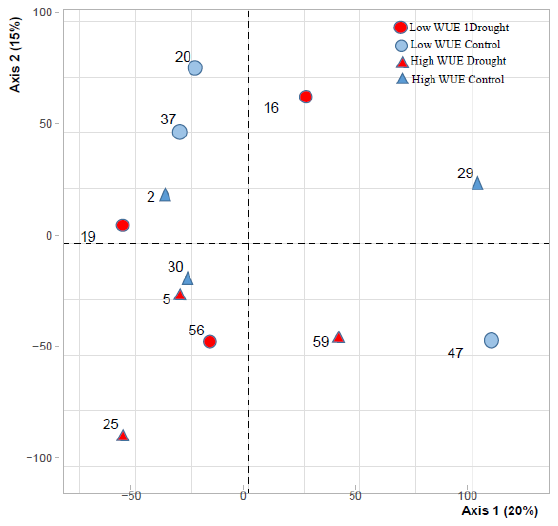
(A)

(B)

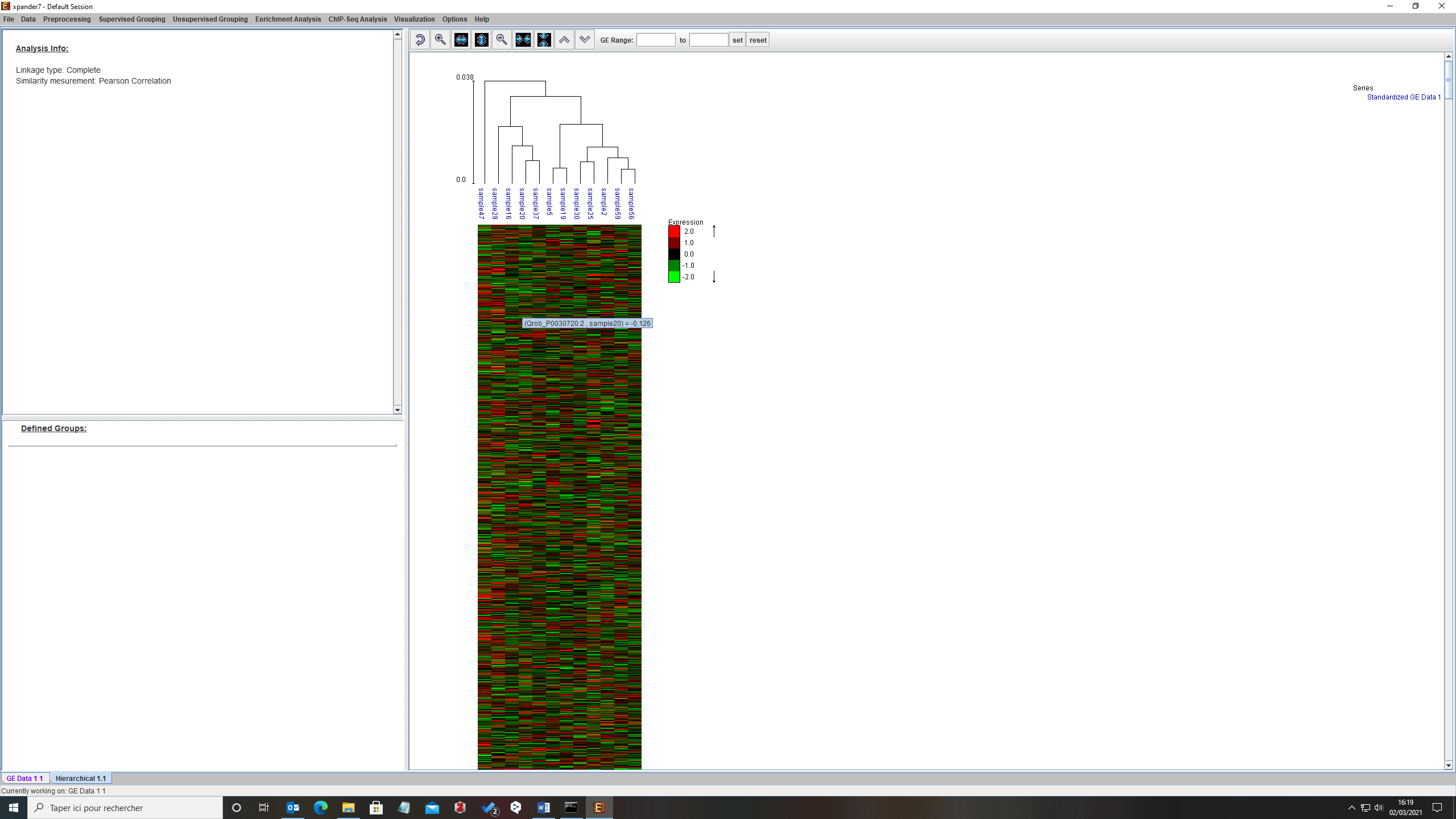

**Low WUE control (47)**

**Low WUE control (20)**

**Low WUE control (37)**

**Low WUE drought (16)**

**Low WUE drought (19)**

**Low WUE drought (56)**

**High WUE control (29)**

**High WUE control (30)**

**High WUE control (2)**

**High WUE drought (5)**

**High WUE drought (25)**

**High WUE drought (59)**
