## Supplementary material for "Molecular plasticity to soil water deficit differs between sessile oak (*Quercus Petraea* (Matt.) Liebl.) high- and low-water use efficiency genotypes": Venn diagram showing the overlap between the three effects analyzed in this study.

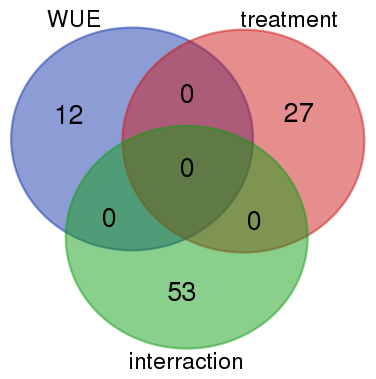
Supplementary. Figure 2 : Venn diagram showing the overlap between the three effects analyzed in this study. We used an adjusted p-value<0.05 to declare a gene differentially expressed.

G*E

E

G
