## Supplementary material for "Molecular plasticity to soil water deficit differs between sessile oak (*Quercus Petraea* (Matt.) Liebl.) high- and low-water use efficiency genotypes": qPCR validation of the candidate genes selected in this study.

Supplementary Figure 3: qPCR validation of the candidate genes regulated by WUE (panel A) or displaying a significant WUE-by-Treatment interaction effect (Cluster#1, panel B, Cluster#2 panel C). Abbreviations correspond to HWUE: High WUE, LWUE: Low WUE, C: Control condition and D: Drought condition. Standard deviations were obtained from the three biological replicates.

**(A)**

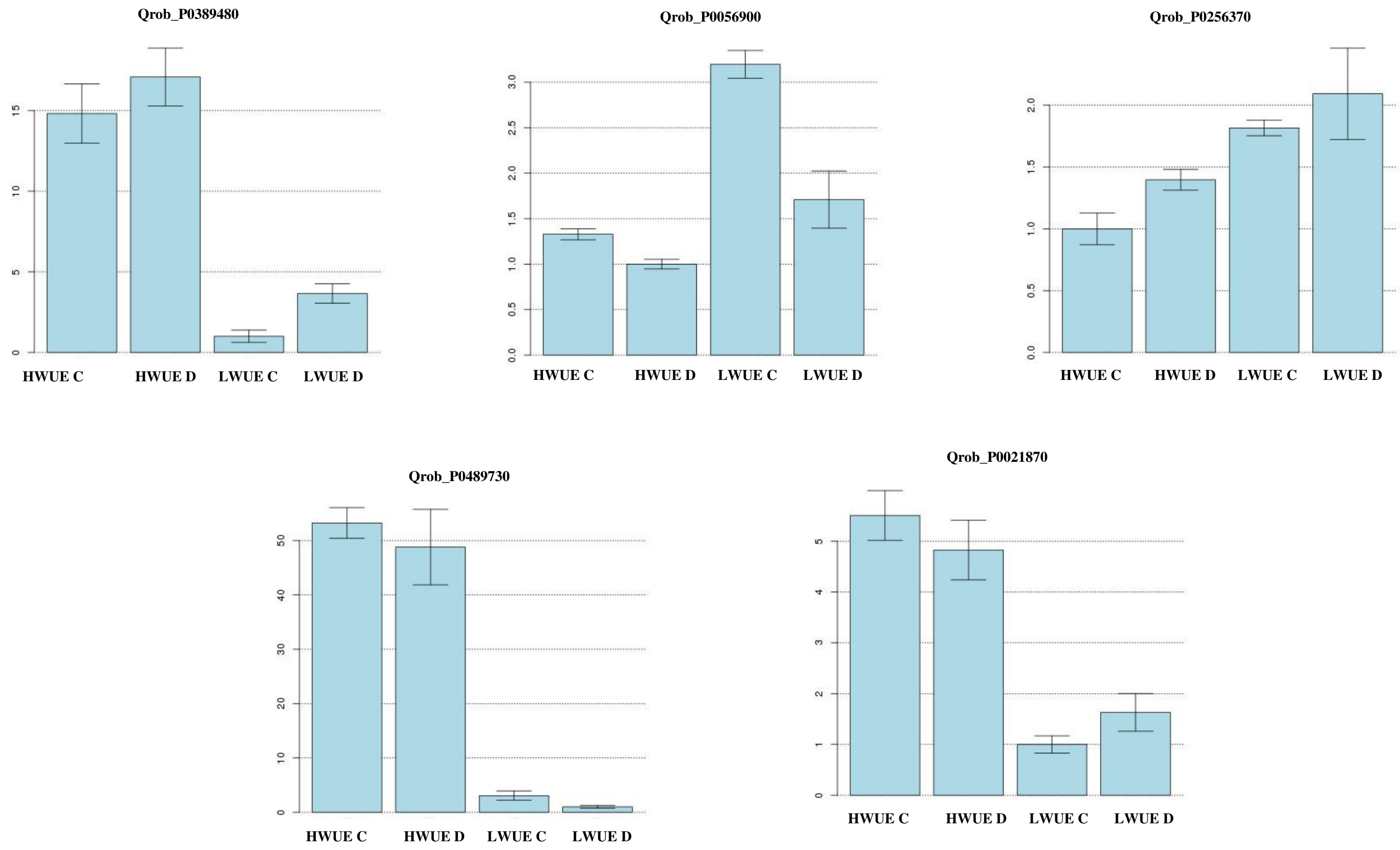

**(B)**

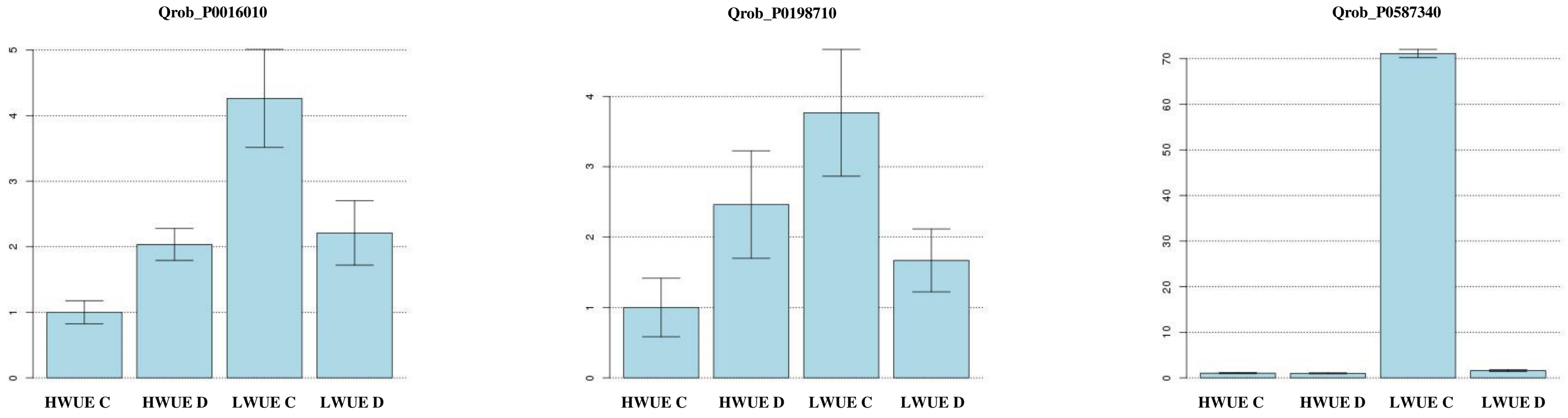
Qrob_P0016010
0
1
2
3
4
5
HWUE C
HWUE D
LWUE C
LWUE D
Qrob_P0198710
0
1
2
3
4
HWUE C
HWUE D
LWUE C
LWUE D
Qrob_P0587340
0
10
20
30
40
50
60
70
HWUE C
HWUE D
LWUE C
LWUE D

(C)

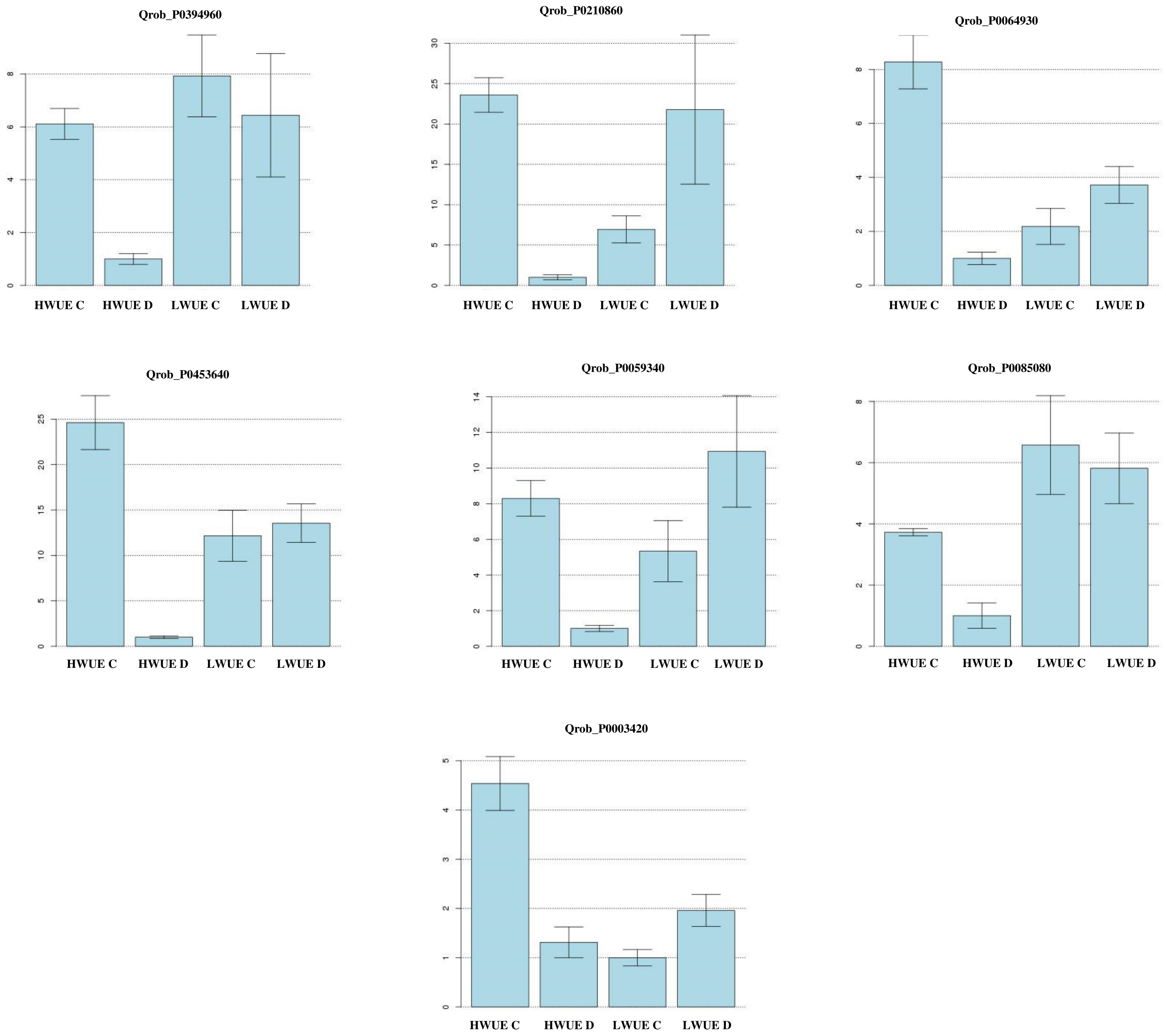

Qrob_P0394960
HWUE C
HWUE D
LWUE C
LWUE D
Qrob_P0210860
HWUE C
HWUE D
LWUE C
LWUE D
Qrob_P0064930
HWUE C
HWUE D
LWUE C
LWUE D
Qrob_P0453640
HWUE C
HWUE D
LWUE C
LWUE D
Qrob_P0059340
HWUE C
HWUE D
LWUE C
LWUE D
Qrob_P0085080
HWUE C
HWUE D
LWUE C
LWUE D
Qrob_P0003420
HWUE C
HWUE D
LWUE C
LWUE D
