## Supplementary material for "Molecular plasticity to soil water deficit differs between sessile oak (*Quercus Petraea* (Matt.) Liebl.) high- and low-water use efficiency genotypes": List of the primer pairs used for qPCR analysis.

**Supplementary Table 2**: List of the primer pairs used for qPCR analysis. Abbreviations: Tm: annealing temperature, HWUE: High WUE, LWUE: Low WUS, CL1: Cluster#1, CL2: Cluster#2, ND: not differentially expressed, NA: not available. Qrob_IDs were retrieved from the oak genome available in Plomion et al. (2018).

.

| Gene ID | Significant effect in  RNAseq | Function | Forward primer(5’3’)  Reverse primer (3’-5’) | Amplicon  size (bp) | Multiband in  agarose gel | Used in  qPCR | PCR  Efficiency | Highest expression in qPCR |
| --- | --- | --- | --- | --- | --- | --- | --- | --- |
| Qrob_P0389480 | WUE | **Ankyrin repeat family protein** | TATTTGGCGGCAGAAGGAGG  CCCGCAGTGAAGGAACCATA | 164 | No | yes | 103 | **HWUE** |
| Qrob_P0056900 | WUE | **NAD(P)-linked oxidoreductase superfamily protein** | CCAGGGGAATATGAGCTGCC CTGGAGGGATCTTGGCTGTG | 175 | No | yes | 108 | **LWUE** |
| Qrob_P0244690 | WUE | **Receptor Like Protein 15** | TCACAATAGCACACCACCCA GCACCATGAAAGGCCTTGTG | 170 | yes | no | NA | **NA** |
| Qrob_P0256370 | WUE | **BEL1-Like Homeodomain 9** | CCACCAACCTACTCACCACC GTACCCAGTAAACGGCCCAA | 169 | No | yes | 95 | **LWUE** |
| Qrob_P0489730 | WUE | **G-type lectin S-receptor-like serine/threonine-protein kinase** | GACACCATCTCTGCACACCA GTCTCTCTGTTTGCCACCCA | 179 | No | yes | 97 | **HWUE** |
| Qrob_P0021870 | WUE | **Leucine-rich repeat receptor-like protein kinase** | GTGGCTCCATTCCTGCTTCT CTGGTGGAATGGAGGCAGAG | 162 | No | yes | 96 | **HWUE** |
| Qrob_P0748930 | WUE*Treatment (CL1) | **aspartyl protease family protein / AP2** | CCCCACCAATAGCCCAACATa ACGCTCGTGATCACAATCCT | 112 | Yes | No | NA | NA |
| Qrob_P0255490 | WUE*Treatment (CL1) | **F-box protein CPR30** | TACACATGCCGCGTCATACAg ACCCATTGCAAGAACCGACT | 116 | Yes | No | NA | NA |
| Qrob_P0016010 | WUE*Treatment (CL1) | **Ribosomal protein S3Ae** | CTCTGTTCCTCACTCGCCAG CGTGAGCACATTCCTCCCTT | 115 | No | Yes | 110 | **CL1** |
| Qrob_P0198710 | WUE*Treatment (CL1) | **Terpene synthase 02** | TCTTGGATGAGACCCTGGCA  AACACCATTGGCATCCTCCC | 123 | No | Yes | 98 | **CL1** |
| Qrob_P0587340 | WUE*Treatment (CL1) | **G-type lectin S-receptor-like serine** | TGATCGGTGGTAGAGAGCCA GCCTATGTCAAACCACCCCA | 121 | No | Yes | 102 | **CL1** |
| Qrob_P0125190 | WUE*Treatment (CL1) | **UDP-Glucosyl Transferase 71C4** | GCTAGTGCGGCATTTCTTGG  GCAGCATACCCATCGTCCTT | 144 | Yes | NO | NA | **NA** |
| Qrob_P0394960 | WUE*Treatment (CL2) | **CBRLK1** | GGAATTGGTCGAGGTCTGCT  GCCATAAGTTCCAACGACCC | 189 | No | Yes | 98 | **CL2** |
| Qrob_P0210860 | WUE*Treatment (CL2) | **PR5-like receptor kinase; GbBRLK5** | TCTTAGTGGGTTTGTGGTGCA GGAGGATTCTTCTGCGGGAG | 155 | No | Yes | 98 | **CL2** |
| Qrob_P0064930 | WUE*Treatment (CL2) | **Spermidine Hydroxycinnamoyl Transferase** | GGCAAAGTTGTCTCGGGGAT GCATCCGTCCTCCCAATCAA | 153 | No | Yes | 98 | **CL2** |
| Qrob_P0453640 | WUE*Treatment (CL2) | **ABC transporter A family** | AACCGACAACTGGCATGGAT TGCAACGAAGCCTACCCTTT | 162 | No | Yes | 98 | **CL2** |
| Qrob_P0273440 | WUE*Treatment (CL2) | **Late embryogenesis abundant (LEA)** | CCATGCCCCTAACTGACGAT TGAATCCGTCACTGTTGGCA | 177 | Yes | No | NA | **NA** |
| Qrob_P0059340 | WUE*Treatment (CL2) | **hypothetical protein; (DUF247)** | GTCGAACCAAGAAAAGCCCG GGTTCGCTTAGTCCTCCACA | 170 | No | Yes | 98 | **CL2** |
| Qrob_P0186810 | WUE*Treatment (CL2) | **WAK2; wall-associated kinase 2** | GAGTGCGAGCGATATCCCAA AGACTTTGGTGTGGGTGGTG | 165 | Yes | No | NA | **NA** |
| Qrob_P0085080 | WUE*Treatment (CL2) | **CNGC1; Cyclic Nucleotide Gated Channel** | CGTCCAAGCGAAGTTGTTGA TGGTCCACAGCACTTCGTTT | 193 | No | Yes | 98 | **CL2** |
| Qrob_P0003420 | WUE*Treatment (CL2) | **SAG101; Senescence Associated Gene 101** | GGCAAATTTCGTGGTGACCC TTGCTACTAGAGGGCCAGGT | 171 | No | Yes | 99 | **CL2** |
| **Control Genes** | | | | | | | | |
| Qrob_P0530610 | NS | **Unknown protein** | GAAGCACCACCCTCACAAGT  GTCTCCTCACAACTCACCGG | 197 | No | Yes | 110 | **NA** |
| Qrob_P0426000 | NS | **Unknown protein** | GCAGAGCTCCAGGACATGATaCAGCAGCAGAGATGAACCCA | 175 | No | Yes | 106 | **NA** |
